# Adversarial Attacks on Protein Language Models

**DOI:** 10.1101/2022.10.24.513465

**Authors:** Ginevra Carbone, Francesca Cuturello, Luca Bortolussi, Alberto Cazzaniga

## Abstract

Deep Learning models for protein structure prediction, such as AlphaFold2, leverage Transformer architectures and their attention mechanism to capture structural and functional properties of amino acid sequences. Despite the high accuracy of predictions, biologically insignificant perturbations of the input sequences, or even single point mutations, can lead to substantially different 3d structures. On the other hand, protein language models are often insensitive to biologically relevant mutations that induce misfolding or dysfunction (e.g. missense mutations). Precisely, predictions of the 3d coordinates do not reveal the structure-disruptive effect of these mutations. Therefore, there is an evident inconsistency between the biological importance of mutations and the resulting change in structural prediction. Inspired by this problem, we introduce the concept of adversarial perturbation of protein sequences in continuous embedding spaces of protein language models. Our method relies on attention scores to detect the most vulnerable amino acid positions in the input sequences. *Adversarial mutations* are biologically diverse from their references and are able to significantly alter the resulting 3d structures.

## 1 Introduction

Understanding how protein sequences fold in a 3d structure has substantial implications for human health, especially in drug design and in the development of disease therapies. SOTA models for structure prediction (e.g. AlphaFold2 [8]) and Transformer based protein Language Models (LMs) [21, 19, 5] are able to recover biological properties of proteins by processing the amino acids in a sequence as words in a sentence. Despite the remarkable capabilities of protein language models, they are often unable to predict misfolding caused by single-point mutations [2]. Moreover, it has been observed that biologically small perturbations in amino acid sequences manage to induce radical changes in the 3d structures [7]. This limitation is presumably due to the scarcity of structure disruptive mutations among the available PDBs [10], which makes protein language models unable to catch all the biological features causing structural variations. We introduce a set of mutations, which we refer to as *adversarial mutations*, whose goal is to: (1) alter a small subset of residues in the original sequences; (2) produce perturbations that are biologically distant from the native sequences. Specifically, we use BLOSUM distance (Section 2) as a metric of biological similarity; (3) induce misfolding with respect to wild-type reference structures. Such mutations could then be used to augment the available training datasets and to make structure prediction LMs more sensitive to disruptive mutations.

## 2 Methodology

We search for biologically plausible perturbations of a few amino acids in the original sequences that are maximally different in the 3d structures. It is natural to interpret this reasoning through the lens of adversarial attacks [6], where the goal is to craft small perturbations of the inputs while inducing the most significant change in predictions. A similar approach is presented in [7], where the authors already propose a notion of “adversarial mutation” on amino acid sequences and use the perturbations to assess the reliability of structure prediction on the RoseTTAFold [1] model. Specifically, they define a robustness measure for structure prediction based on the computation of the inverse RMSD between original and perturbed structures and use it to produce adversarial sequences. The fundamental novelty in our approach is that the adversarial perturbations do not require direct knowledge of the 3d coordinates, but only leverage the hidden representations provided by language models.

In the large space of all possible 3d structures, only a small portion contains biologically meaningful proteins. Moreover, high confident scores in structure prediction using AlphaFold2 model do not necessarily guarantee the plausibility of the native sequences [11]. Generative models [12, 9, 20, 11] tackle this problem by learning the data distribution in the space of sequences. Hence, they catch new evolutionary dependencies between the amino acids and generate new samples from the learned distribution. Our method does not directly rely on a generative architecture, but rather explores the space of hidden representations while guaranteeing that a set of desired conditions are met. In what follows, we discuss the approach behind our choices of target positions and mutant residues.

### Target positions

Several recent works show that Transformer language models are able to recover functional properties of protein sequences [18, 26, 21]. Vig et al. [26], in particular, observe that attention scores capture structural information and that most of the attention is directed to binding sites, i.e. amino acids that are far apart in the sequence but close in the 3d structure. We use attention scores to identify token positions having the highest impact on the surrounding context (i.e. on the remaining amino acids) and use them as target positions for adversarial mutations. Let *x* = (*x*_1_, …, *x*_*N*_) ∈ 𝒜^*N*^ be a protein sequence, where 𝒜 is the 25-character alphabet of amino acids^1^ and *N* ∈ ℕ is the length of the sequence. Given a fixed number of token substitutions *n < N*, each *attention head h* ∈ {1, …, *H*} in a Transformer LM computes a set of attention scores *A*_*h,l*_(*x*) for each layer *l*∈ {1, …, *L*} in the architecture. The resulting target tokens are the first *n* positional indexes maximizing the Euclidean norm of the average attention across all layers and heads, i.e. the first *n* values in arg sort_*i*=1,…,*N*_ ‖𝔼_*h,l*_[*A*_*h,l*_(*x*_*i*_)]‖_2_.

### Target residues

Once target token indexes are fixed, we use Block Substitutions Matrices (BLOSUM) [25] to assess the set of allowed substitutions of residues at each position. BLOSUM matrix contains integer similarity scores between all couples of residues in a sequence. In particular, scores in the popular BLOSUM62 matrix are based on replacement frequencies observed in known alignments with less than 62% sequence similarity [7]. The selection of amino acids at a given position is restricted to residues with non-null BLOSUM scores, i.e. those with non-null frequency in the reference alignments, to avoid biologically meaningless substitutions. We also use BLOSUM62 matrix at evaluation time to assess the biological similarity between perturbed and original sequences. The choice of target residues is performed through a search over all plausible token substitutions that satisfy a desired “adversarial” property. Given a fixed set of target positions *I* in a wild-type sequence *x*, we propose multiple attack strategies to craft an adversarial perturbation 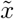, built upon different hidden representations of the original sequence but with the common goal of causing a significant structural change:

- Let *z* and 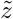 respectively denote the continuous embeddings of *x* and 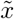 in the first embedding space, i.e. the positional embeddings. **Maximum distance perturbations** indicate token substitutions 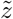 that maximize the L1 distance from *z* in the first embedding space, i.e.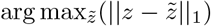;
- Protein LMs are trained to solve a *masked prediction task*, meaning that part of the input sequence is masked at random positions, and the model has to predict missing residues from the surrounding context. Therefore, given single candidate residue at a target position *i*, the LM outputs a *pseudo-likelihood* score 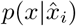, denoting an approximate likelihood of the full sequence with the chosen residue, where 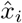 is the sequence masked at position *i*. We define a loss function 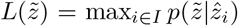 that penalizes the highest pseudo-likelihood score attributed to the first embedding of an adversarial sequence 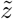 and use it to build a perturbation inspired by classical gradient-based attacks in continuous spaces [6]. **Maximum cosine similarity perturbations** search for residues that maximize the cosine similarity w.r.t. the loss gradient direction in the first embedding space, i.e. they build perturbations in the direction of greatest change in the loss function. More precisely, given a gradient-based attack 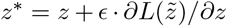 in the first embedding space, they search for a 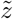 that maximizes cos similarity 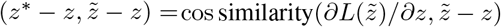. Notice that this definition does not depend on the intensity *ϵ* of the attack;
- Protein LMs provide a reduced representation of the 3d structure, known as *contact map*, which consists of a heatmap of estimated distances between all residue pairs in the 3d structure [27]. **Maximum contact map distance perturbations** maximize the L2 distance between original and perturbed contact maps: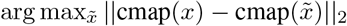;
- Additionally, we introduce **maximum entropy perturbations** for MSA Transformer only. Given an input sequence *x* with an associated MSA and a set of amino-acid substitutions 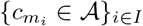 at target sites *I*, we use residue substitution frequencies 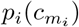 in the MSA to compute the entropy of that substitution. Then, we search for a perturbation that maximizes the entropy: 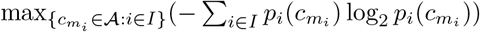.

### Evaluation metrics

Since protein LMs for structure prediction could be insensitive to single point mutations [2], we do not only rely on evaluation metrics on the structure [16], but also examine several evaluation metrics on continuous embeddings of amino acid sequences. The first natural evaluation metric for sequence similarity is the L1 distance between original and perturbed embeddings in the first layer of a protein LM, 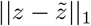, providing a preliminary geometric interpretation of the distribution of continuous representations in the first embedding space. We point out that the choice of the first embedding space to evaluate distances provides a natural baseline for comparison (i.e. maximum distance perturbations), but it would be interesting to extend this analysis to multiple layers of hidden representations. Secondly, we leverage BLOSUM62 matrix to compute a biological sequence similarity measure known as BLOSUM distance: 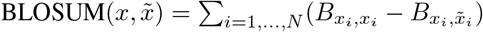, where 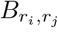 is the entry associated to a couple of residues (*r*_*i*_, *r*_*j*_) in BLOSUM62 matrix. BLOSUM distance is zero when 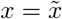 and positive otherwise. Another fundamental information provided by protein LMs is the predicted contact map, thus we also compute the L2 distance between contact maps of original and perturbed sequences. It is usual practice to examine upper submatrices in a contact map and look at the distances between long-range contacts. Therefore, we use an index *k* ∈ ℕ_*>*0_ to denote the diagonal index of an upper triangular submatrix in a full contact map (*k* = 0), i.e. we select contacts that are at least *k* positions apart, and compute a range of distances 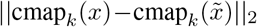 as *k* increases.

On the structural side, let *s* and 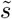 respectively denote original and perturbed 3d structures. We evaluate their similarity by means of both local and global similarity measures to capture the relative orientation of the deviations as well as the global superimposition between the structures. Local Distance Difference Test (LDDT) [13] is a local score that measures the percentage of preserved distances between all pairs of atoms in the target structure closer in space than a predefined cutoff. In particular, it computes the mean fraction of preserved distances using four different thresholds (0.5, 1, 2, 4 *Å*). Then, we use two popular global scores: Root-Mean-Square-Deviation (RMSD) computes the average Euclidean distance between matching atoms in the two structures; TM-score, instead, computes the degree of match between corresponding C*α* atoms, scaled by a length-dependent distance parameter. Detailed definitions of the structural measures are provided by [16].

## 3 Experimental results

Our experiments^2^ involve two Transformer models trained on Uniref50 database [24], namely ESM1b [21] and MSA Transformer [19]. ESM-1b takes as inputs protein sequences and computes pairwise attention scores between all couples of amino acids. The set of attention scores *R*_*h,l*_(*x*) ∈ ℝ^*N*^ from an attention head *h* in the *l*-th layer is called *row attention*, where *x* is the input sequence and *N* is the length of the sequence. The resulting attention score used to recover target positions is *A*_*h*_(*x*) := 𝔼 _*l*_[*R*_*h,l*_(*x*)]. MSA Transformer, instead, works on MSAs [3], i.e. on sets of aligned protein sequences. In this setting, self-attention mechanism also computes *column attention* scores *C*_*h,l*_(*x*) ∈ ℝ^*N*^ from the input MSA and we select positions based on a weighted sum of the two scores: *A*_*h*_(*x*) := *γ* 𝔼 _*l*_[*R*_*h,l*_(*x*)] + (1 − *γ*) 𝔼 _*l*_[*C*_*h,l*_(*x*)]. In the experiments, we weight the two contributions equally by setting *γ* = 0.5. We use the hhfilter method from HH-suite tool [23] to select a subset of most diverse sequences in the alignment by means of a sequence similarity score based on the degree of homology among sequences. We build a filtered MSA for each selected sequence to be used as an input for MSA Transformer. Then, we craft the set of adversarial perturbations presented in Section 2.

First, we build adversarial mutations on protein families PF00533 and PF00627, both containing examples of structure disruptive mutations not detected by AlphaFold2 model [2]. For the sake of brevity, we only report the experiments performed on domain PF00627 in the main text, where we use MSA Transformer model to build 3 sites mutations on 100 native sequences with input MSAs of depth 100. We observe a similar experimental behaviour on domain PF00533 and include the additional experiments in the Appendix. Next, we compare adversarial perturbations to those provided by ProTherm database [15], containing more than 4*k* single-point mutations associated with high changes in stability. Protein stability denotes the capability of a protein to retain the native conformation under a stress condition (e.g. change in temperature or pressure). Since high changes in stability are often associated with disease-causing mutations and unfolding, we want to analyze their statistical properties against those of adversarial mutations. We focus on ProTherm values of *change in free energy* ΔΔ*G* to select the most stabilizing (highest ΔΔ*G*) and destabilizing (lowest ΔΔ*G*) mutations. Specifically, we set a threshold of 1*kcal/mol* on the minimum absolute value of ΔΔ*G*. In this case, we identify the reference sequences belonging to Pfam families and build the associated filtered MSA accordingly.

### Impact of adversarial mutations on wild-type sequences

We analyze the effect of adversarial perturbations using the evaluation metrics described in Section 2. First of all, Figure 1 shows how adversarial perturbations are overall less likely compared to the residues selected by masked predictions, suggesting that our method detects rare substitutions at fixed mutant sites. This is an important argument in favour of the generation of new and diverse disruptive mutations. We compare adversarial perturbations to the ones obtained from all the discarded plausible token substitutions at the chosen target positions, which we refer to as “other” in Figures 2a and 3a. Figure 2 reports the L1 distances 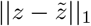 between original and perturbed first layer embeddings, while Figure 3 shows BLOSUM distances between original and perturbed sequences. Adversarial perturbations significantly depart from the other plausible perturbations both in terms of embedding distance (Figure 2a) and BLOSUM distance (Figure 3a), approaching perturbations at maximum embedding distance in the first case. Adversarial embeddings are at least as far from the reference as the most stabilizing and destabilizing embeddings in ProTherm (Figure 2b), while BLOSUM distances are comparable (Figure 3b). Moreover, Jha et al. [7] observed that larger BLOSUM distances between original and perturbed sequences lead to higher RMSD in predicted structures, therefore we expect adversarial perturbations to produce a significant structural change. We stress that adversarial mutant positions for ProTherm are those that maximize the attention scores and, in most cases, differ from ProTherm mutation sites. Nonetheless, adversarial sequences are able to significantly alter embedding distances, BLOSUM distances and contact maps distances (see the Appendix for additional results), suggesting that our attention-based selection method catches new relevant mutant positions.

**Figure 1:**
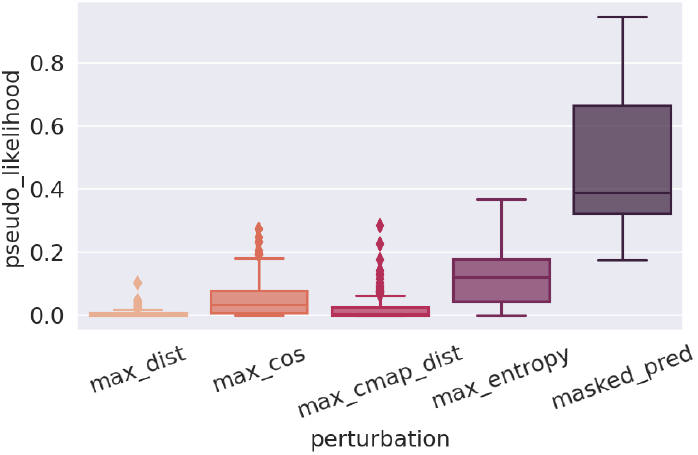
Pseudo-likelihood of adversarial (columns 1-4) and masked prediction (column 5) mutations at target token indexes. Values refer to 3 sites mutations obtained from MSA Transformer on 100 sequences from domain PF00627.

**Figure 2:**
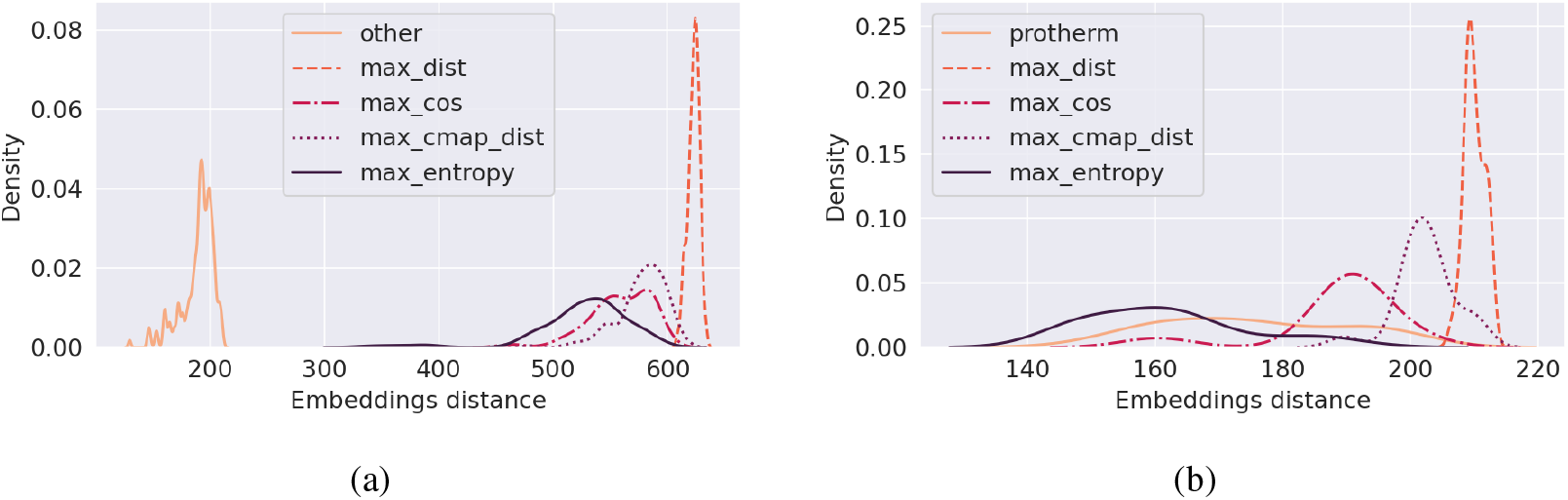
L1 distances 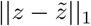 between the embeddings of original and perturbed sequences in the first embedding space. Perturbations are computed using MSA Transformer model. In (a) we report mutations of 3 sites on domain PF00627, while Figure (b) shows single mutations on ProTherm database. “Other” refers to adversarial perturbations obtained from all the discarded plausible token substitutions at the chosen target positions.

**Figure 3:**
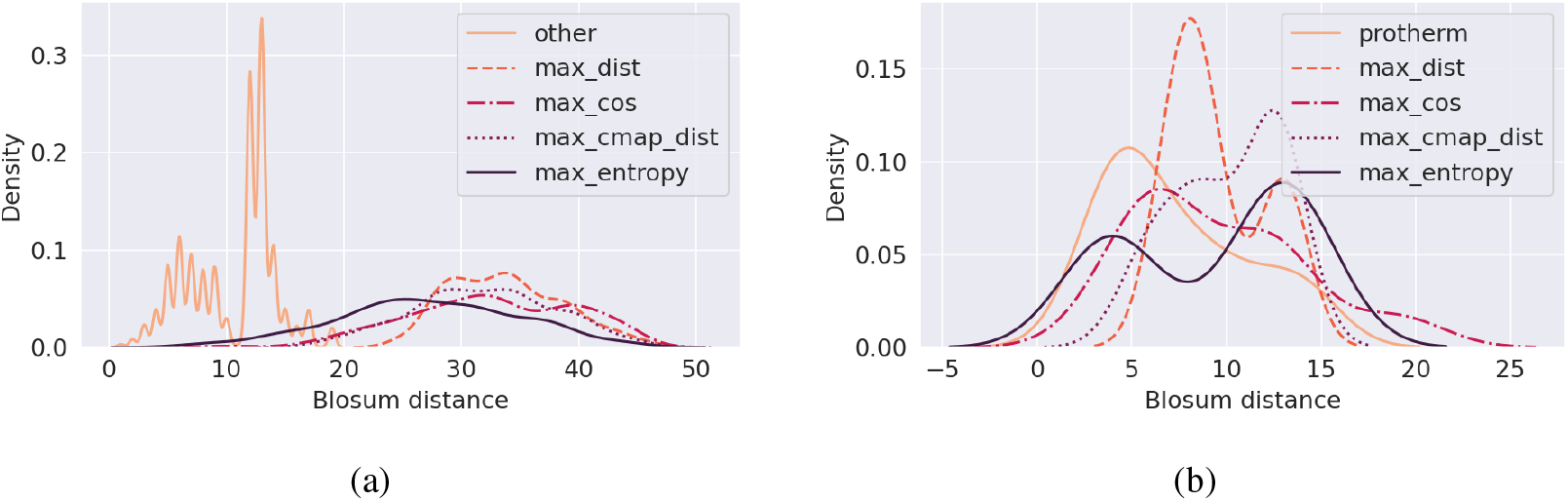
Blosum distances BLOSUM 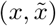between original and perturbed sequences. Perturbations are computed using MSA Transformer model. In (a) we report mutations of 3 sites on domain PF00627, while (b) shows single mutations on ProTherm database. “Other” refers to adversarial perturbations obtained from all the discarded plausible token substitutions at the chosen target positions.

### Predicted adversarial structures are substantially altered and highly confident

We analyze the effect of adversarial perturbations on 3d structures predicted by ColabFold [14], an extension of AlphaFold2 model. First, we select the 100 most diverse sequences from domain PF00627 using hhfilter method, as explained at the beginning of this chapter. Since our goal is to generate adversarial mutations that are able to “fool” structure prediction models, the latter should be highly confident in predictions. Therefore, we rely on the pLDDT confidence score for structure prediction provided by AlphaFold2 model, and among the 100 original sequences we select the ones whose average (over residues) pLDDT is greater than 80%, for a total of 37 sequences. Then, we build the adversarial perturbations, predict their 3d structures and compare them to the original predicted structures. Figure 4 reports LDDT, TM-score and RMSD (presented in Section 2) between original and perturbed 3d structures on among the selected sequences. Figure 11 in the Appendix reports the resulting adversarial pLDDT scores, as well as another accuracy measure for structure prediction, namely the pTM-score. Adversarial mutations exhibit high confidence in structure prediction and produce structures that are significantly distant from their references in terms of global distance. In particular, low TM-scores and high RMSD scores, consistently across the several attack techniques, indicate that adversarial structures deviate from the original folding. As a matter of comparison, it is important to observe that two structures with 50% sequence identity align within approximately 1*Å* RMSD [4], and two proteins with even 40% sequence identity and at least 35 aligned residues align within approximately 2.5*Å*[22]. Higher LDDT scores in Figure 4 instead show that local atomic interactions in the original structure happen to be preserved in adversarial structures, especially for maximum entropy perturbations.

**Figure 4:**
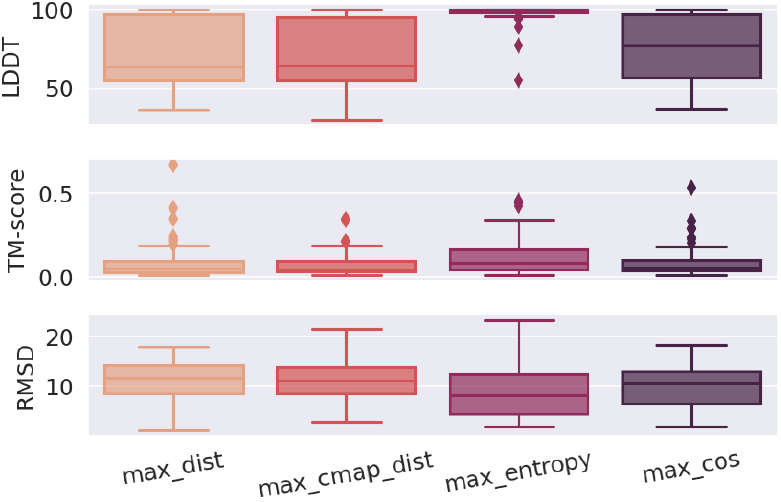
LDDT, TM, and RMSD scores between original and perturbed structures for 3 sites mutations on 100 sequences from domain PF00627. Adversarial perturbations are computed by MSA Transformer model, while structure predictions are performed in ColabFold.

## 4 Conclusions and future directions

Adversarial perturbations on Protein Language Models produce substantial changes according to several geometric, biological, and structural evaluation scores compared to the reference sequences. Additionally, they introduce a new efficient attention-based method for the selection of target positions in the reference sequences. Nonetheless, we point out the presence of some limitations, which we plan to address in future versions of this work. Precisely, we intend to: (1) propose a more extensive empirical evaluation on several protein families and include structure prediction on databases of dysfunctional mutations, (2) fine-tune protein LMs on adversarial sequences and test their sensitivity to a set of known missense mutations. We believe that adversarial perturbations could help to improve the sensitivity of protein LMs to dysfunctional mutations, therefore opening a new path for a deeper understanding of the connection between protein stability and variations in the 3d structures.

## Acknowledgments and Disclosure of Funding

The authors acknowledge AREA Science Park supercomputing platform ORFEO made available for conducting the research reported in this paper, and the technical support of the staff of the Laboratory of Data Engineering. F.C. was supported by the grant PNR “FAIR-by-design”. A.C. was supported by the ARGO funding program.

## Appendix

Here we report some additional experiments performed on domain PF00533 using ESM-1b model (Figures 6, 7a, 7b, 8a) and MSA Transformer model (Figures 12, 13); on domain PF00627 using ESM-1b model (Figure 8b) and MSA Transformer model (Figures 5, 11); on ProTherm database, using ESM-1b model (Figures 9a, 10a, 10b) and MSA Transformer model (Figures 10a, 10b).

**Figure 5:**
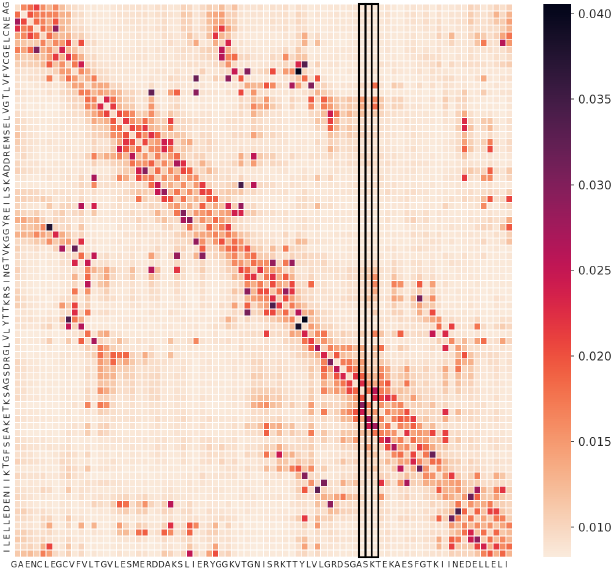
Average attention scores over all attention heads for an input sequence from domain PF00627. Scores are computed by MSA Transformer.

**Figure 6:**
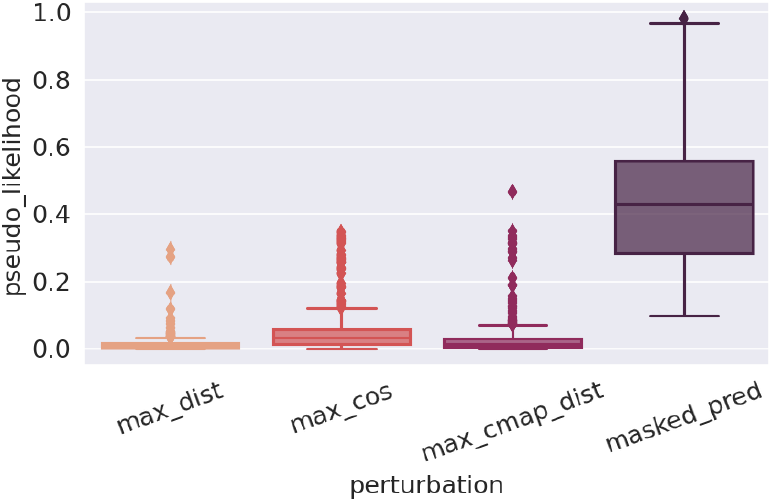
Pseudo-likelihood of adversarial (columns 1-3) and masked prediction (column 4) mutations at target token indexes. Values refer to 3 sites mutations obtained from ESM-1b on 100 sequences from domain PF00533.

**Figure 7:**
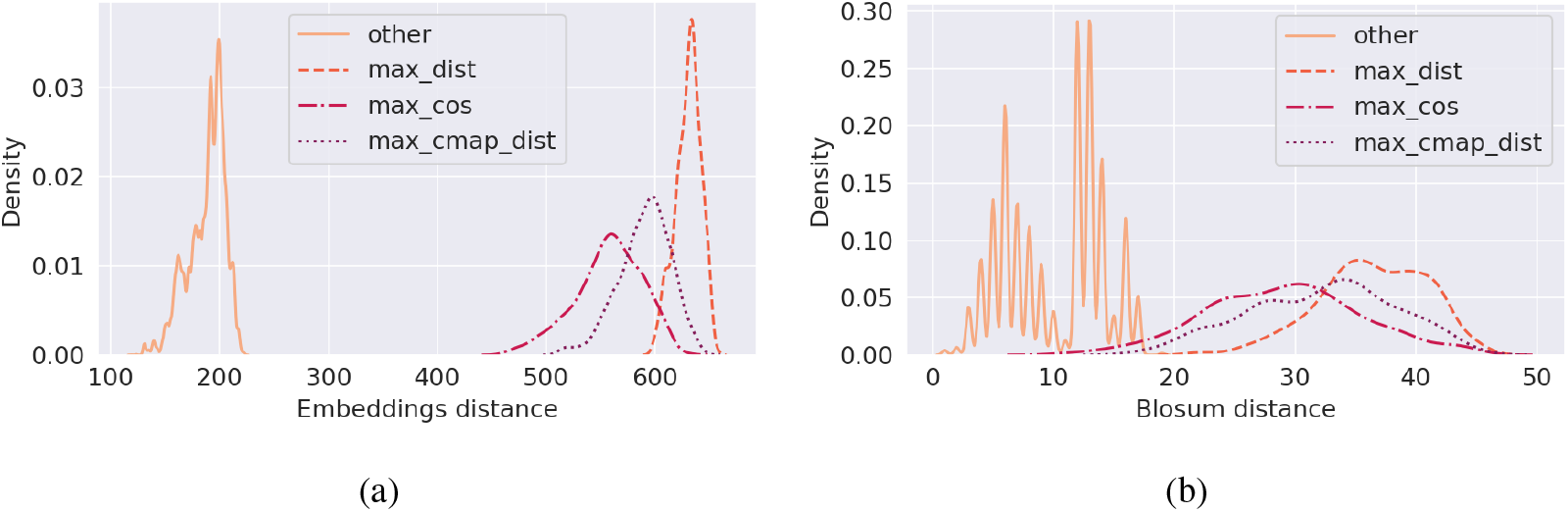
L1 distances 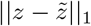 between the embeddings of original and perturbed sequences in the first embedding space (a) and Blosum distances BLOSUM 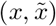between original and perturbed sequences. Adversarial mutations on 3 sites are computed using ESM-1b model on domain PF00533. “Other” refers to adversarial perturbations obtained from all the discarded plausible token substitutions at the chosen target positions.

**Figure 8:**
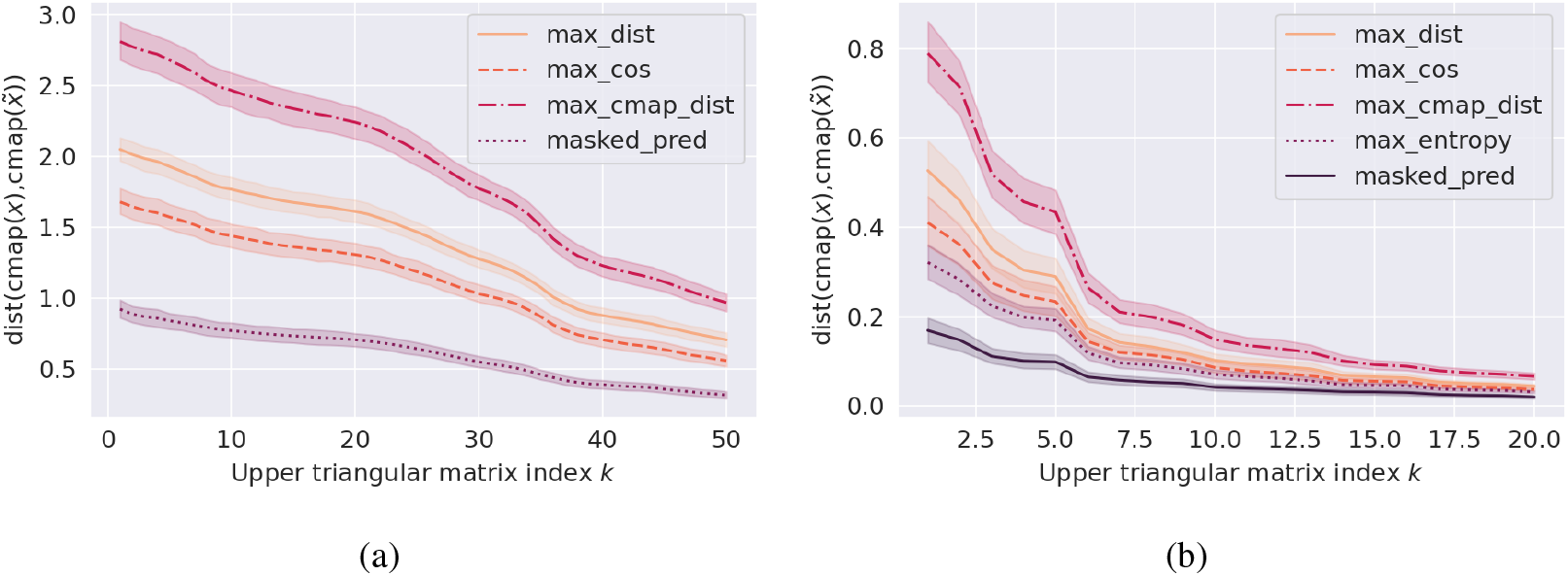
Distances between upper submatrices of contact maps of original and perturbed sequences, as the index *k* of upper triangular submatrices increases. Adversarial perturbations at 3 sites are computed by ESM-1b model 100 sequences from domain PF00533 in (a) and by MSA Transformer model on 100 sequences from domain PF00627 in (b).

**Figure 9:**
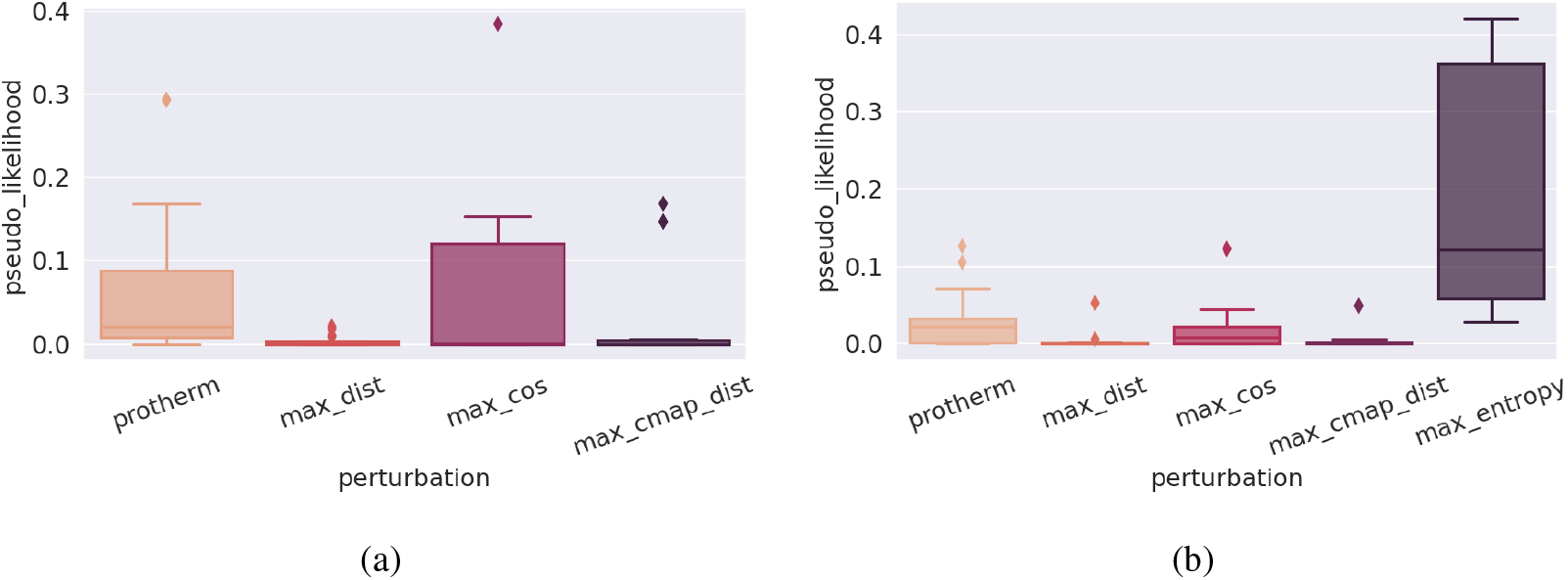
Pseudo-likelihood of ProTherm (first column) and adversarial (other columns) mutations at target token indexes. Values refer to single residue mutations obtained on ProTherm database using ESM-1b model in (a) and MSA Transformer model in (b).

**Figure 10:**
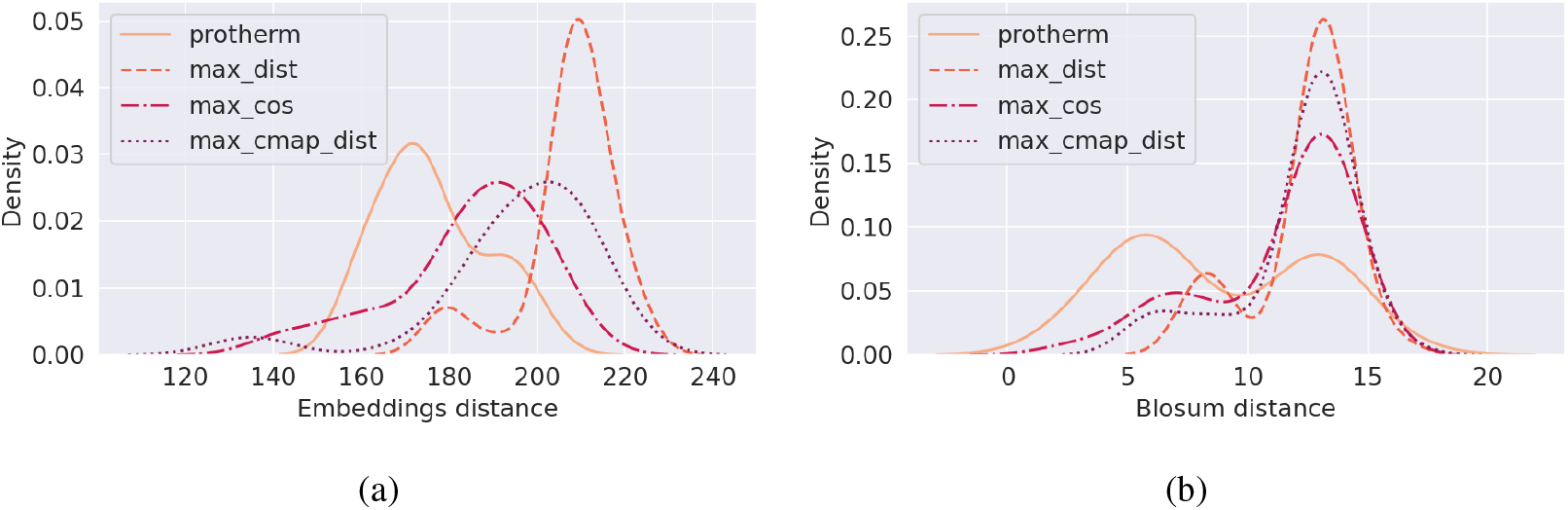
L1 distances 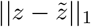 between the embeddings of original and perturbed sequences in the first embedding space (a) and Blosum distances BLOSUM 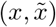 between original and perturbed sequences. Adversarial mutations on 3 sites are computed using ESM-1b model on ProTherm database.

**Figure 11:**
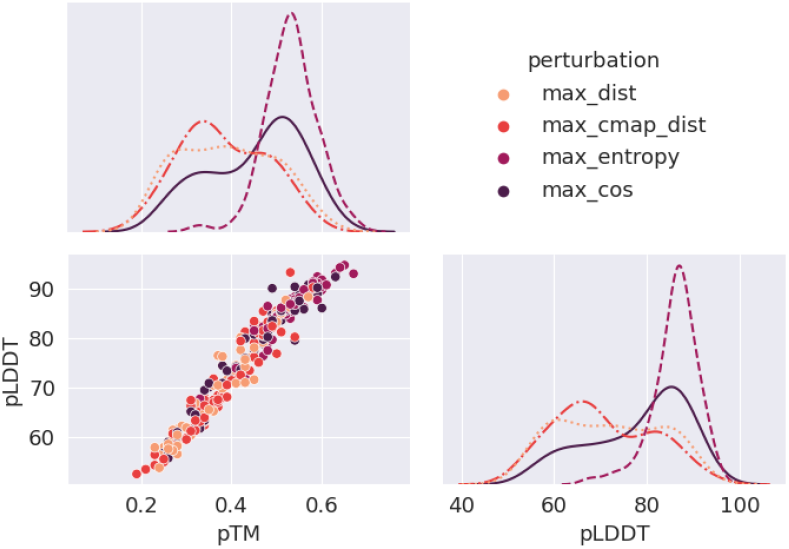
Confidence scores for structure prediction performed in ColabFold on adversarial sequences from PF00627 such that pLDDT *>* 80 on the original structures.

**Figure 12:**
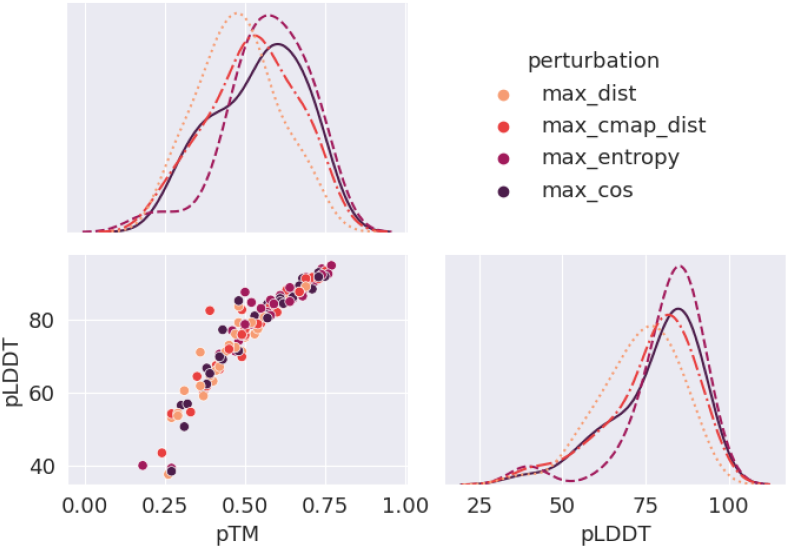
Confidence scores for structure prediction performed in ColabFold on adversarial sequences from PF00533 such that pLDDT *>* 80 on the original structures.

**Figure 13:**
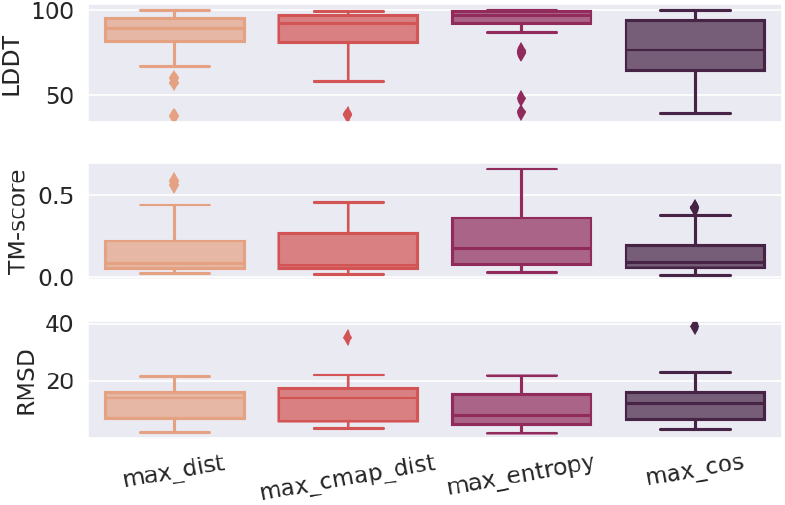
LDDT, TM and RMSD scores between original and perturbed structures for 3 sites mutations on 100 sequences from domain PF00533. Adversarial perturbations are computed by MSA Transformer model, while structure predictions are performed in ColabFold.

## Footnotes

1 20 characters for the standard amino acids and 5 characters for non-standard or unknown amino acids [17].

2 Code is available at https://github.com/ginevracoal/adversarial-protein-sequences.

